# BAX and BAK expression triggers multiple IRE1 signaling outputs under ER stress

**DOI:** 10.1101/2024.03.26.586784

**Authors:** Claudia Sepulveda, Mateus Milani, Vania Morales, Giovanni Tamburinni, Nicolas Montes, Claudio Hetz

## Abstract

Adaptation to endoplasmic reticulum (ER) stress depends on the activation of the unfolded protein response (UPR) stress sensor inositol-requiring enzyme 1 alpha (IRE1). IRE1 is a central ER stress sensors, that signals through the activation of its RNase domain to catalyze the splicing the mRNA encoding the transcription factor X-box binding protein 1 (XBP1), resulting on the expression of a stable and active transcription factor termed XBP1s. The kinetics and amplitude of IRE1 signaling are regulated by different posttranslational modifications and the physical interaction of different factors. Early studies demonstrated that the expression of the proapoptotic proteins BAX and BAX enhance UPR signaling. However, the possible effects on the RNase activity were not defined. Here we provide preliminary evidence indicating that BAX and BAK deficiency increases the in 10 folds the threshold of ER stress to induce XBP1 mRNA splicing, and the upregulation of its target genes. In addition, the degradation of RIDD substrates was strongly reduced in BAX and BAK null cells. BAX and BAK double deficiency also attenuated the levels of IRE1 phosphorylation under mild ER stress. These results reinforce previous findings indicating that proapoptotic BAX and BAK have alternative functions at the ER regulating the UPR.

## Introduction

Sustaining proteostasis is fundamental for organismal health, and its deregulation contributes to a series of chronic disease in addition to normal aging^1,2^. The endoplasmic reticulum (ER) is a central node of the proteostasis network involved in protein folding and secretion, in addition to operating as a central site for calcium storage and lipid synthesis. Multiple physiological and pathological conditions favor the accumulation of misfolded proteins in the ER lumen, resulting in a cellular state referred to as ER stress^3^. In fact, chronic ER stress is emerging as a relevant factor contributing to various diseases, including metabolic syndromes, cancer, diabetes, inflammatory diseases, and neurodegeneration^4^. The unfolded protein response (UPR) is the main adaptive mechanism to cope with ER stress and restore proteostasis^5^. Inositol-requiring enzyme 1 alpha (IRE1) is a type I ER transmembrane protein with a serine and threonine protein kinase and endoribonuclease activity, that upon activation, catalyzes the splicing of the mRNA encoding X-box binding protein 1 (XBP1), leading to the expression of a potent transcription factor termed XBP1s (for the <u>s</u>pliced form)^5^. XBP1s regulates a cluster of genes involved in different aspects of the secretory pathway, including protein folding, ER-associated degradation (ERAD), protein quality control, among others^6,7^. The RNase activity of IRE1 also degrades selected mRNAs and microRNAs through a process known as regulated IRE1-dependent decay (RIDD), contributing to inflammation, DNA damage, apoptosis, and other processes^8^.

There is increasing evidence that the signaling behavior of UPR transducers IRE1, PERK and ATF6 are modulated by the binding to specific factors^3^. Thus, the threshold of ER stress that triggers the UPR is determined by specific interactomes, which may influence the adaptive capacity of a cell and the susceptibility to undergo apoptosis under ER stress. Multiple laboratories have identified positive regulators of IRE1α signalling that function by controlling IRE1α dimerization, oligomerization, phosphorylation and dephosphorylation, impacting the amplitude and kinetics of the signalling response (reviewed in ^3^). Our lab reported that several members of the BCL-2 family physically associate with IRE1 to enhance the amplitude of UPR signaling, including proapoptotic BAX and BAK and upstream regulators such as BIM and PUMA^9,10^. In contrast, the antiapoptotic protein BAX-inhibitor 1 BI-1 negatively regulates IRE1, impacting the UPR attenuation process^11^. More than 30 proteins have been identified to bind and regulate IRE1 function in different cellular systems, highlighting non-muscle myosin heavy chain IIB protein, the tyrosine-protein kinase ABL1, the collagen carrier HSP47, the core component of the translocon machinery Sec61PKA, among others (reviewed in ^3^). Thus, IRE1 signaling is a highly regulated process involving distinct checkpoints, defining the threshold to trigger an adaptive UPR or transit into a terminal cell death program.

Early studies showed that BAX and BAK double deficiency (DKO) reduces the expression of XBP1s protein in cells and animals exposed to experimental ER stress, associated with a physical interaction^9^. However, many assays were not available at that time to monitor the IRE1 activation process and its RNase activity. Here we investigated the possible impact of BAX and BAK expression on the endoribonuclease activity of IRE1 was not studied. Here we provide preliminary evidence indicating that the threshold to induce XBP1 mRNA slicing is reduced in BAX and BAK DKO cells. These effects were associated with a reduction in the upregulation of XBP1s target genes, in addition to the down regulation of classical RIDD targets. The phosphorylation of IRE1 was also attenuated as measured using PhosTag assays and non-denaturing electrophoresis. These preliminary results confirm the impact of BAX and BAK expression on IRE1 signaling.

## Results

To define the possible impact of BAX and BAK expression on the activity of IRE1, we performed a dose response experiment in wild type (WT) and BAX and BAK DKO murine embryonic fibroblast (MEFs). We exposed cells to different concentrations of tunicamycin (Tm, 0.05-1 μg/ml) and measured XBP1 mRNA splicing after 4 h using conventional RT-PCR. Our results indicated that BAX and BAK DKO cells were 10-folds less sensitive to process of XBP1 mRNA splicing (Figure 1a). These results were confirmed using PCR primers that selectively amplify the processed XBP1s mRNA (Figure 1b). In agreement with these results, the expression of the XBP1s protein as reduced in BAX and BAK DKO cells (Figure 1c).

**Figure 1.**
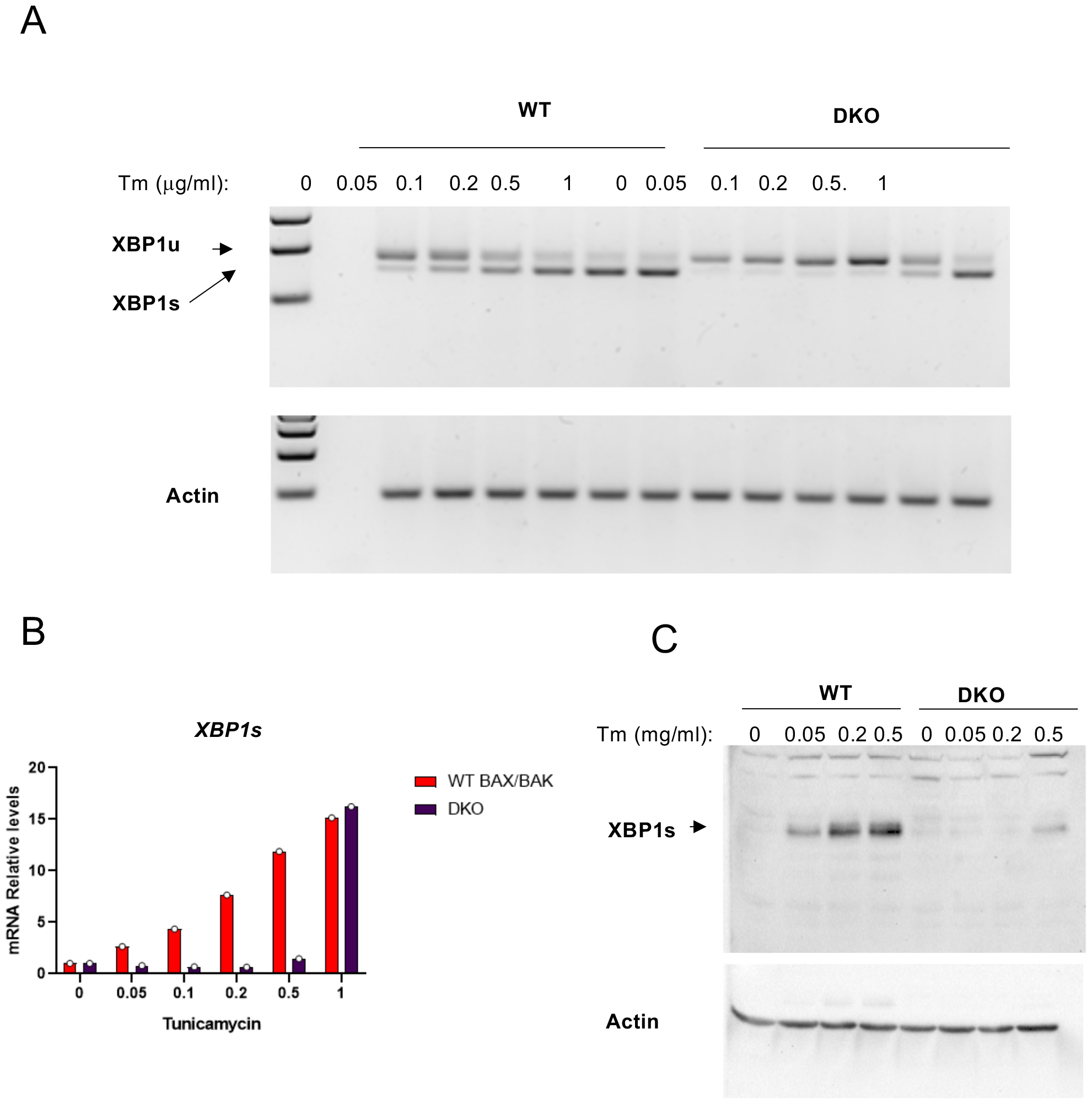
BAX and BAK double deficiency reduces XBP1 mRNA splicing under ER stress. **(A)**To monitor IRE1 RNase activity, BAX and BAK WT and double knockout (DKO) MEFs were treated with indicated concentrations of tunicamycin (Tm) for 4 h and then RNA was extracted and analyzed by RT-PCR to detect both the unspliced XBP1u and the XBP1s forms. Actin was amplified as control. **(B)** We also confirmed the effects of BAX and BAK double deficiency on XBP1 mRNA splicing using real time PCR primers that specifically amplify the XBP1s form. **(C)** BAX and BAK WT and DKO MEFs were treated with indicated concentrations of tunicamycin (Tm) for 6 h and then analyzed by Western blot to measure XBP1s expression.

We also monitored the consequences of BAX and BAK double deficiency on IRE1-reppedent transcriptional responses under ER stress. The upregulation of XBP1s target genes *Erdj4, Edem1* and *Sec61* were reduced in BAX and BAK DKO cells treated with Tm (Figure 2a). In addition to catalyze XBP1 mRNA splicing, IRE1 degrades a subset of mRNAs through a process termed RIDD. We monitored the levels of *BlosC1*, a canonical RIDD target, under ER stress, and observed that BAX and BAK double deficiency reduced RIDD activity, suggesting the regulation of distinct signaling outputs (Figure 2b). Similar results were observed when the mRNA levels of the RIDD substrates *Spark* and *Col6a* were monitored by quantitative PCR (Figure 2c). Finally, we monitored other signaling outputs controlled by IRE1. Under ER stress, IRE1 binds TRAF2 to recruit JNK and induce its activation^12^. Phosphorylation of JNK was reduced under ER stress in BAX and BAK DKO cells. However, normal activation was observed in cells treated with TNF alpha or UV exposure (Figure 3). BAX and BAK controls ER calcium content to regulate cell death, a phenomena that can ge genetically corrected by ovexpressing SERCA. WE used this system to assess the activation of the UPR in BAX and BAK DKO cells. Overexpression of SERCA did not increase the levels of XBP1 mRNA splicing and UPR signaling (suplementary figure 1).

**Figure 2.**
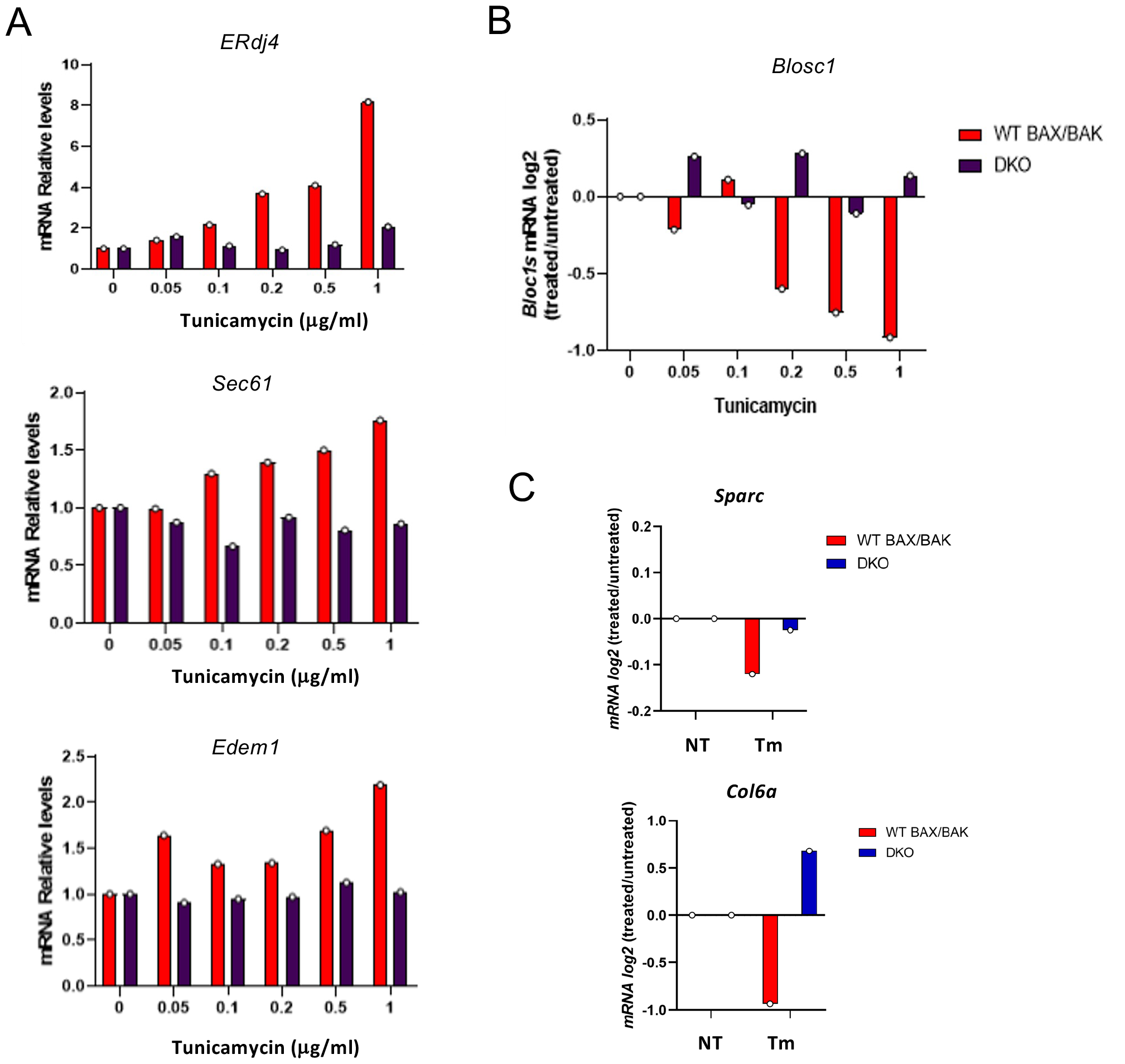
Reduced expression of XBP1s-target genes and RIDD substrate degradation in BAX and BAK double knockout cells. **(A)** BAX and BAK WT and DKO MEFs were treated with indicated concentrations of tunicamycin for 4 h and then RNA was extracted and analyzed by real time PCR to measure the canonical XBP1s target genes *ERdj4, Sec61* and *Edem1*. **(B)** In addition to process the XBP1 mRNA, IRE1 degrades certain mRNAs through RIDD. We measured the decay of the canonical RIDD substrate *Blosc1* under ER stress in the same samples. **(C)** In addition, the mRNA levels of alternative RIDD substrates *Sparc* and *Col6a* were assessed in cells treated with 1 ug/ml of tunicamycin.

**Figure 3.**
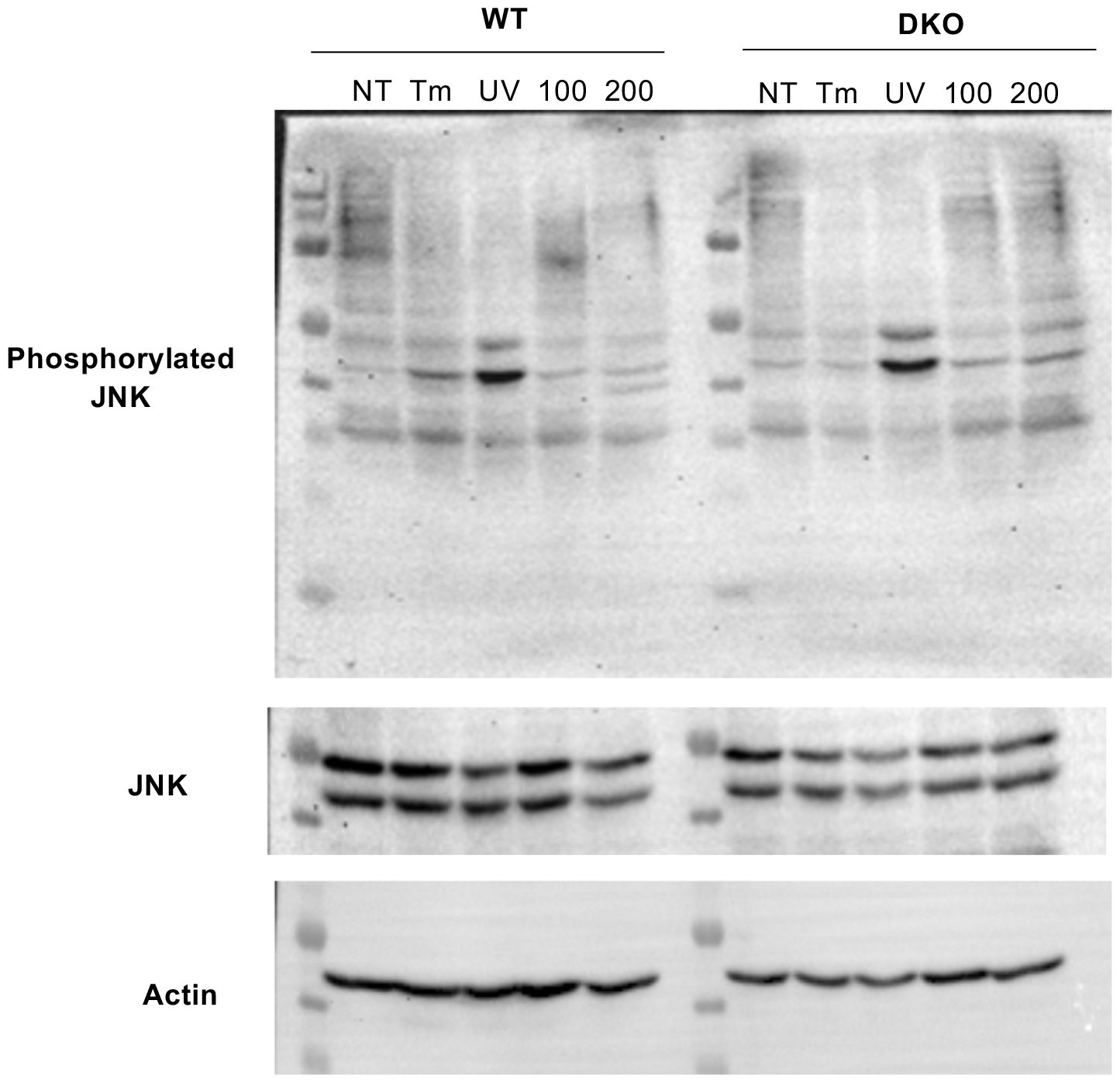
BAX and BAK DKO cell show reduced JNK phosphorylation under ER stress. BAX and BAK WT and DKO cells were treated incubated in cell culture media with 1% FBS and then treated with 1 μg/mg tunicamycin (Tm), 100 or 200 ng/ml TNF (100, 200), or exposed to UV light (UV) or left untreated (NT) and phosphorylated JNK was monitored by western blot analysis.

Activation of IRE1 involves its oligomerization into large clusters. We used human HEK293 T cells expressing an inducible form of IRE1 fused to GFP. Then we knocked down BAX, BAK or both together (Figure 4A). After 24 h, IRE1-GFP expression was induced with doxicicline and then 24h later we stimulated cells with Tm. Analisis of IRE1-GFP foci formation indicated a reguction when BAR and BAK were knocked down (Figure 4B and C). IRE1 activation involves the auto-phosphorylation of the kinase domain, resulting on a conformational change that engages its RNase domain. We monitored the phosphorylation of IRE1 using a PhosTag assay. Dose response experiments indicated that BAX and BAK doble deficiency increased the threshold of ER stress required for IRE1 phosphorylation (Figure 5). Similar results were obtained using non-denaturing gels (Figure 5).

**Figure 4.**
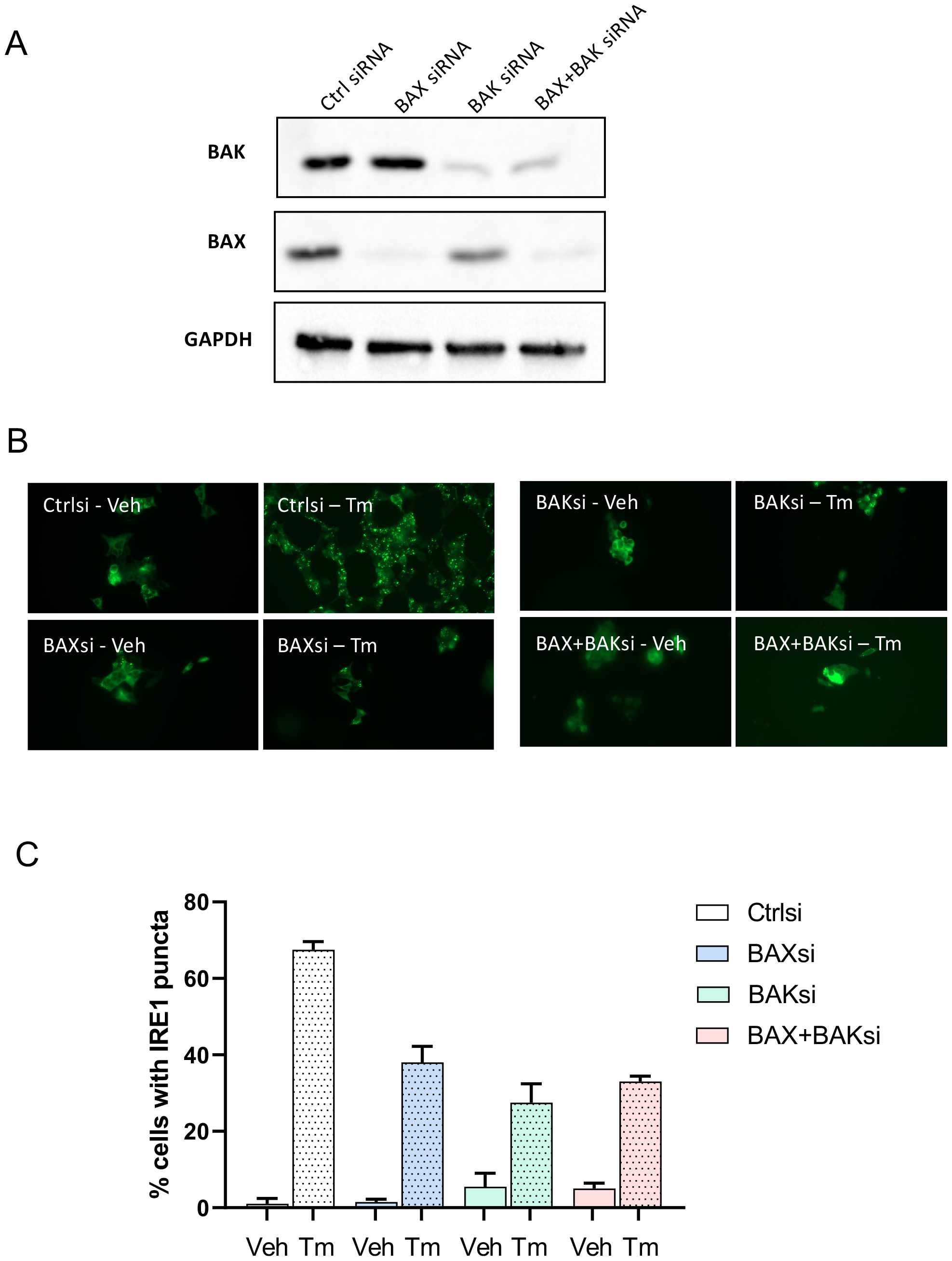
Analisis of IRE1 oligomerization (clustering) in Bax and BAK fedidcient cells. 293T cells expressing Ire1-GFP on an inducible manner were transfected witgh siRNAs for BAX, BAK, or both together or a control siRNA. Then, after 28 h cells were treated with 1 uM dox to induce IRE1 expression and 24 h later treated with 300 ng/ml of the ER stress agent tunicamycin (Tm). The percentage of cells containing IRE1-GFP foci was quantified by fluorescent microscopy at 4 h.

**Figure 5.**
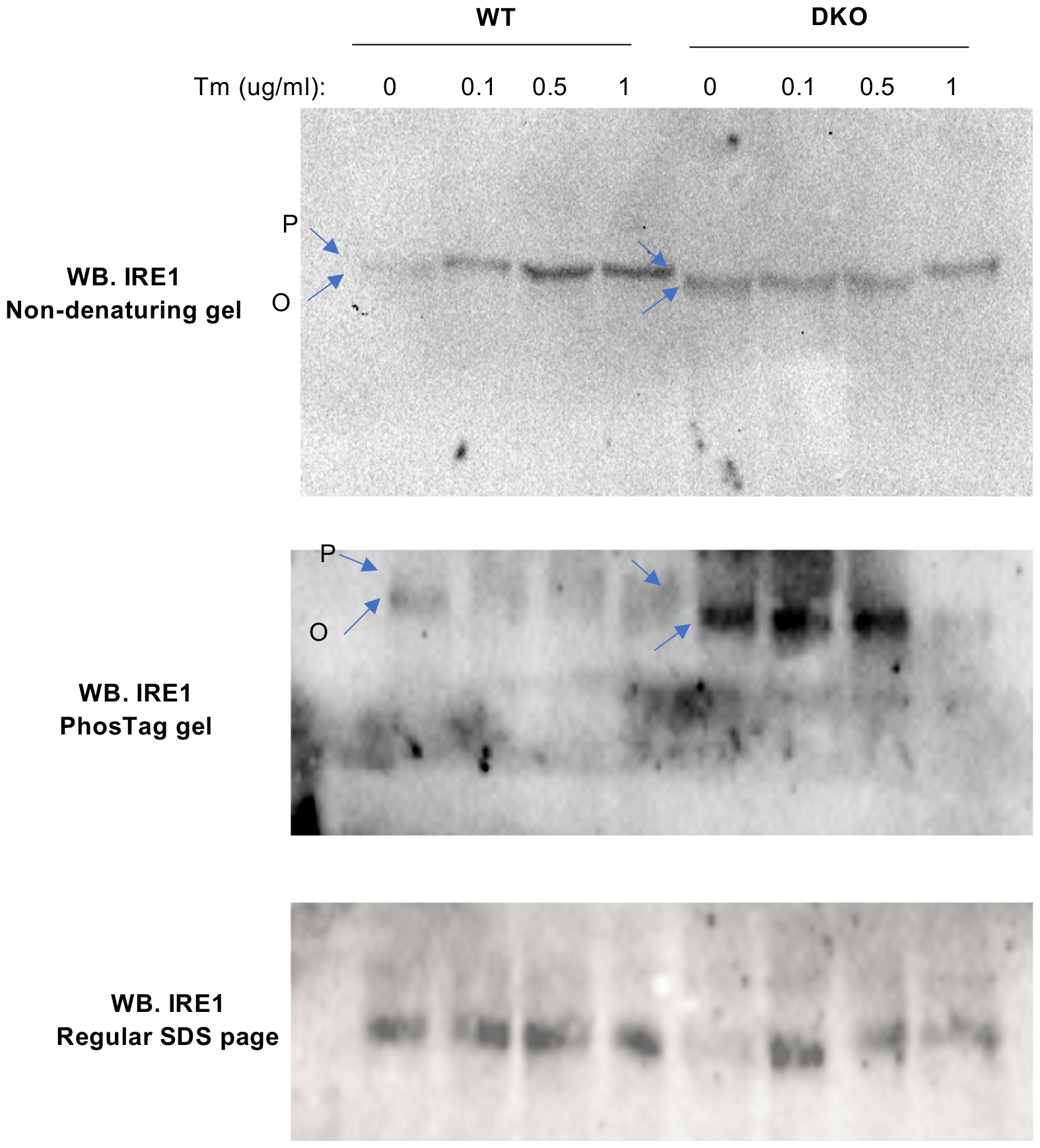
Effects of BAX and BAK deficiency on IRE1 phosphorylation. BAX and BAK WT and DKO cells were treated with indicated concentrations of tunicamycin (Tm) for 6 h. Protein extracts were analyzed in 3 different types of western blot analysis: non denaturing gels, PhosTag gels and normal electrophoresis. A shift in the molecular weight of IRE1 was detected (P).

## Conclusions

IRE1 initiates the most conserved signaling pathway of the UPR, determining the recovery of proteostasis under ER stress. IRE1 function has been implicated on a variety of diseases, and the use of small molecules to target the activity of IRE1 (RNase and kinase) have demonstrated important protective effects in various preclinical models of disease^13^. At the molecular level, a complex network of regulatory checkpoints tightly control IRE1 signaling behavior. The concept of the UPRosome (or IRE1 signalosome) was proposed to illustrate the idea that the activity of IRE1 is regulated by cofactors, in addition to crosstalk with other stress signaling pathways mediated by the assembly of adapter proteins and signaling mediators^14^.

Our group have uncovered different IRE1 interactors using unbiased approaches (yeast two hybrids and IP-mass spectrometry analysis) identifying PUMA/BIM as novel regulators of IRE1 upstream of BAX and BAK^10^, in addition to the chaperone and collagen carrier Hsp47^15^, or the actin regulator Filamin A^16^. We also showed that cABL interacts with IRE1 to control RIDD^17^, and or with the IP3R to regulate calcium transfer to the mitochondria and bioenergetics^18^. These examples illustrate the highly dynamic nature of the IRE1 interactome and the vast consequences to UPR regulation and the crosstalk with multiple biological processes. Here we have provided additional preliminary evidence confirming a regulatory role of BAX and BAK on the UPR. These results will be part of a full study to determine the biochemical aspects involved in the interaction bteween BAX/BAK and IRE1.

## Materials and methods

### Cell lines

MEF cells used here were described in ^9^, and maintained in Dulbecco’s modified Eagles medium supplemented with 5% fetal bovine serum, non-essential amino acids. The pMSCV-Hygro retrovirus vector expressing IRE1α-HA was previously described^10^. IRE1α contains two tandem HA sequences at the C-terminal domain and a precision enzyme site before the HA tag. COS-1 cells were maintained under standard tissue culture conditions using 10% fetal bovine serum in Dulbecco’s modified Eagles medium (DMEM) (Sigma). HEK cells were maintained in DMEM supplemented with 5% fetal bovine serum.

#### RNA isolation, RT-PCR, and real-time PCR

Total RNA was prepared from cells and tissues using Trizol (Invitrogen, Carlsbad, CA, USA), and cDNA was synthesized with SuperScript III (Invitrogen) using random primers p(dN)6 (Roche). Quantitative real-time PCR reactions employing SYBRgreen fluorescent reagent and/or EvaGreen™ were performed in the Stratagene Mx3000P system (Agilent Technologies, Santa Clara, CA 95051, United States). The relative amounts of mRNAs were calculated from the values of comparative threshold cycle by using Actin as a control and Rpl19 for RIDD. All methods for the *Xbp1* mRNA splicing assay, RIDD and the assessment of XBP1s-target genes used here were previously described^10,11,15^. Real time PCR primers are described in KEY RESOURSCES TABLE.

#### IRE1α oligomerization assay

TREX cells expressing IRE1α-3F6HGFP WT were obtained from Dr. Peter Walter at UCSF. TREX cells plated and treated with doxycycline (500 ng/mL for 24 h). Cells were treated with Tm and fixed with 4% paraformaldehyde for 30 min. Nuclei were stained with Hoechst dye. Coverslips were mounted with Fluoromount G onto slides and visualized by confocal microscopy (Fluoview FV1000). The number and size of IRE1α foci was quantified using segmentation and particle analysis of Image J software.

#### Immunoblot analysis and phostag gels

Immunoblot analysis was performed using standard conditions (Rojas-Rivera et al., 2012). The following antibodies and dilutions were used: Anti-β-actin (1:3000; 5125, Cell Signaling), anti-HA (1:2000; 901514, Biolegend) anti-IRE1alpha (1:1000; 3294, Cell Signaling), anti-Phospho-SAPK/JNK (1:1000; 4668, Cell Signaling) anti-SAPK/JNK (1:1000; 9252 Cell Signaling). Detection of the phosphorylated IRE1α form was performed using the Phostag™ assay loading 15 μg of total protein onto 4% SDS-PAGE minigels containing 80 μM of Phostag™ in the presence of 25 mM MnCl2.

---

**Supplementary Figure S1.**
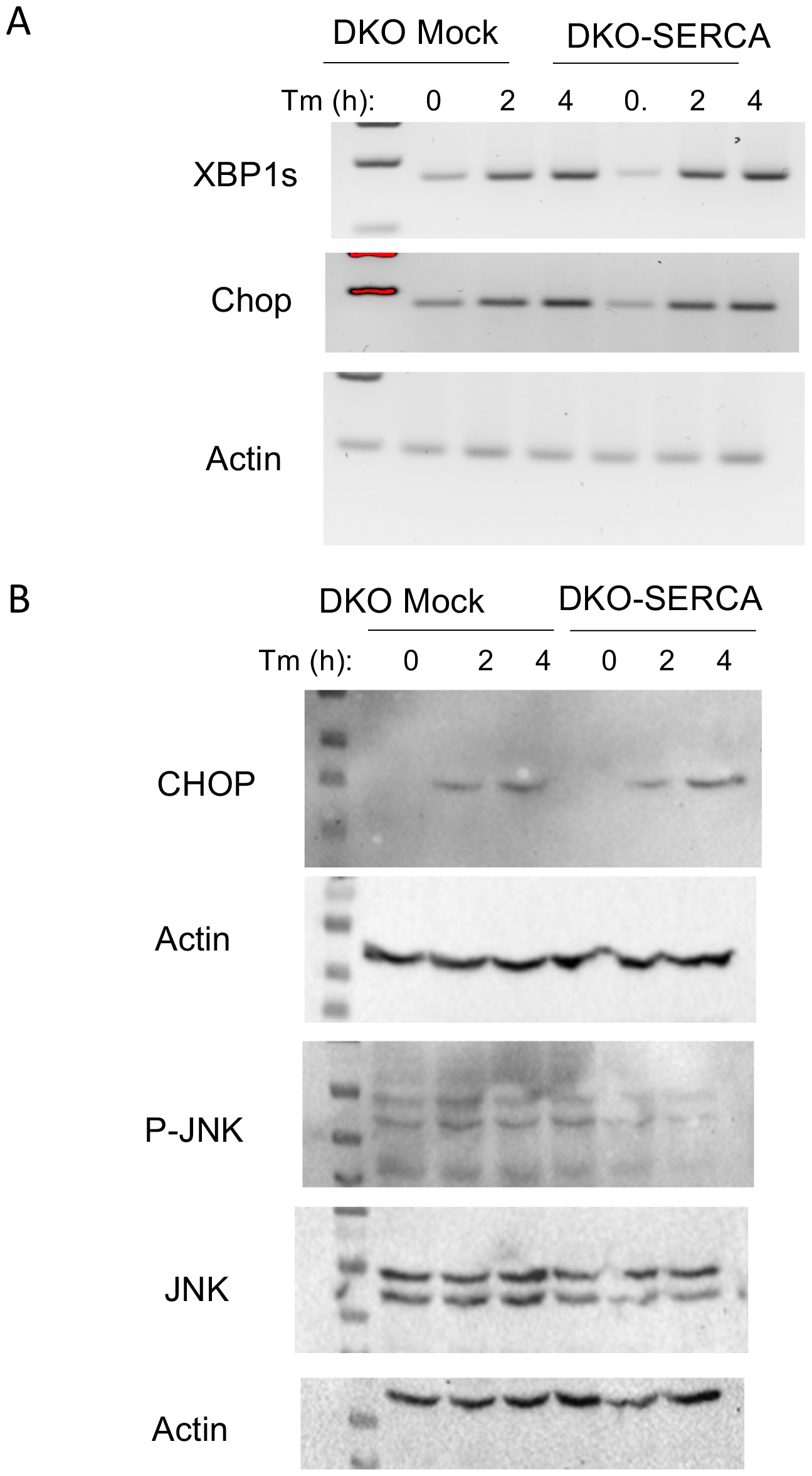
Normal UPR activation in BAX and BAK DKO cells overexpressing SERCA. **(A)** BAX and BAK DKO cells were engineered to overexpress SERCA or empty vector (Mock) to restore the normal ER calcium phenotype (cells from Scorrano, Oakes et al., 2003 Science). Cells were treated with tunicamycin for 2 and 4 h or left untreated. XBP1s and Chop mRNA levels were monitored by semiquantitative RT-PCR. **(B)** In parallel CHOP and phospho-JNK were monitored by western blot analysis.

